# Novel infection by *Mucor hiemalis* kills *Caenorhabditis* hosts through intestinal perforation

**DOI:** 10.1101/2025.06.04.657490

**Authors:** Jay Ni, Jessica N Sowa

## Abstract

The nematode *Caenorhabditis elegans* has emerged as a popular model system to investigate cell biology and host-pathogen interactions. Presently, *C. elegans* is studied as a natural host of intracellular pathogens such as microsporidia and Orsay virus along with extracellular bacterial and fungal pathogens. The use of *C. elegans* as a model in host-pathogen research is limited by the number of naturally occurring pathogens to the organism. Through a sampling project to identify new pathogens of *C. elegans*, we identified the fungus *Mucor hiemalis* as a pathogen of *Caenorhabditis* species. We observed the fungus in the intestinal lumen of wild-caught *Caenorhabditis briggsae,* and co-culturing the wild-caught species with infection reporter *C. elegans* confirmed infection by *M. hiemalis*. This study characterizes the fungal infection by *M. hiemalis* in *Caenorhabditis* nematodes. Fluorescence microscopy with fungal staining revealed the life cycle of *M. hiemalis* within multiple *Caenorhabditis* species at varying growth stages. We observed the killing of nematodes by *M. hiemalis* via intestinal perforation and assessed its’ host range through a series of lifespan assays. We investigated the food preference of *C. elegans* and determined that nematodes show a preference towards food that contains *M. hiemalis* spores. Lastly, we evaluated common *C. elegans* transcriptional immune responses and found that *M. hiemalis* does not induce genes associated with the intracellular pathogen response or other responses seen with previously studied bacterial and fungal pathogens. Characterization of this fungal infection in *Caenorhabditis* nematodes will provide new insights into the biology of pathogenic fungi and host immune responses.

## Introduction

Fungi are a major cause of fungal disease and infection in humans with an annual burden of over 1 billion globally (Bongomin et al., 2017). *Aspergillus, Candida, Cryptococcus, Histoplasma,* and *Mucor* species are the main fungal pathogens responsible for most serious fungal diseases (Bongomin et al., 2017). Fungi cause superficial, mucosal, and invasive infections in tissues with a wide range of severity (Bongomin et al., 2017). Fungal infections in immunocompromised individuals can be deadly and are responsible for over 1.5 million deaths globally per year (Rayens & Norris, 2022). Fungal disease is a pressing area of concern as incidence increases due to a range of global issues including climate change and antibiotic resistance (Bongomin et al., 2017). Fungal infections also further complicate patients with other chronic and immunosuppressive diseases such as HIV/AIDS, tuberculosis, diabetes, asthma, and cancers (Bongomin, 2020; Denning et al., 2006; Rayens et al., 2021; Rayens & Norris, 2022; Rodrigues et al., 2019; Sipsas & Kontoyiannis, 2012). The diverse array of fungal pathogens and their complex relationship with host immune systems highlights a need for further research to understand host-pathogen interactions and identify potential therapeutic targets.

The nematode *Caenorhabditis elegans* is a powerful and genetically tractable invertebrate system to model pathogenic microbes and their relationship with hosts. In the wild, the typical *C. elegans* diet consists of commensal bacteria from decaying organic matter; however they also encounter a variety of microbes which are pathogenic (Shtonda & Avery, 2006). *C. elegans* is a natural host to a range of pathogens including bacteria, fungi, microsporidia, and Orsay virus (Darby, 2005; Félix et al., 2011; Troemel et al., 2008). Research using *C. elegans* as a model for host-pathogen interactions has resulted in significant discoveries in microbial pathogenesis and conserved host immune system responses (Marsh & May, 2012).

*C. elegans* lack an adaptive immune system and do not have any known cellular immunity or specialized immune cells (Tran & Luallen, 2024). *C. elegans* also lack homologs of most canonical mammalian pathogen recognition receptors that identify specific pathogens and their associated damage. However, *C. elegans* does have innate epithelial cell immunity that responds to the presence of pathogenic microbes and their virulence factors, and several evolutionarily conserved immunity pathways have been characterized in *C. elegans*. The p38 mitogen-activated protein kinase pathway is a signaling cascade that responds to infections by *Staphylococcus aureus* or *P. aeruginosa* to provide defense (Irazoqui et al., 2010; Kim et al., 2002). The conserved *C. elegans* transforming growth factor β and DAF-2/insulin signaling pathways provide regulation of innate immunity signaling responses and the secretion of antimicrobial peptides (Zugasti & Ewbank, 2009). The conserved G-protein-coupled receptor DCAR-1in *C. elegans* responds to physical damaging of the epidermal tissue in response to some fungal pathogens (Zugasti et al., 2014). Further, studies of *C. elegans* and its naturally occurring intracellular pathogens have led to the discovery of an immunity signaling cascade comparable to mammalian type-I interferon responses called the intracellular pathogen response (IPR) (Reddy et al., 2017). The IPR consists of eighty genes that are upregulated in response to intracellular infections caused by microsporidia *N. parisii* and Orsay virus (Reddy et al., 2017).

*C. elegans* have been used to model human fungal infections including *Candida, Cryptococcus, Histoplasma,* and *Aspergillus* species to advance understanding of fungal pathogenesis and host immune system responses. *Candida albicans* infects the *C. elegans* intestinal tract via ingestion and causes death by hyphal proliferation (Pukkila-Worley et al., 2011). The PMK-1/p38 MAPK pathway was found to promote resistance to *C. albicans* infection, but the mechanism has yet to be characterized (Pukkila-Worley et al., 2011). *Cryptococcus neoformans* infection has also been modeled in *C. elegans*. The capsule of *C. neoformans* is toxic to *C. elegans* when ingested. Several fungal virulence factors were identified using *C. elegans* as a model (Mylonakis et al., 2002). Insulin/insulin-like growth factor 1 and DAF-16 pathways were observed to provide defense against *Cryptococcus* infection in *C. elegans* (Kitisin et al., 2022). The killing of *C. elegans* by *Aspergillus fumigatus* has also been used to model fungal pathogenicity, but the *C. elegans* immune response to this fungal pathogen has yet to be discovered (Okoli & Bignell, 2015). *Drechmeria coniospora* is a natural pathogen of *C. elegans* and has been isolated in wild populations. *D. coniospora* is not known to induce human infection, but the infection in *C. elegans* has been characterized (Félix & Duveau, 2012). *D. coniospora* attach to the cuticle of nematodes and penetrate the protective layer to cause infection (Félix & Duveau, 2012; Jansson, 1994). An induction of antimicrobial peptide genes related to the conserved p38-MAPK cascade was observed in response to the epidermal wounding caused by *Drechmeria* (Pujol et al., 2008). The relatively limited number of natural fungal pathogens of *C. elegans* limits host-pathogen research, particularly given the current rise of fungal threats to human health.

We isolated the fungus *Mucor hiemalis* from a wild sample of *Caenorhabditis briggsae* through the Nematode Hunters project, which is designed to discover novel natural pathogens of wild nematodes. While testing for an IPR response with reporter *C. elegans*, we observed that *M. hiemalis* can infect other *Caenorhabditis* species including *C. elegans*.

*M. hiemalis b*elongs to the class Zygomycetes, order Mucorales, and genus *Mucor* (Prabhu & Patel, 2004). Fungi from the order Mucorales are opportunistic fungal pathogens that are causative agents of mucormycosis. Mucormycosis infections in immunocompromised hosts are often severe, while infections in immunocompetent hosts are rare (Hernández & Buckley, 2025). In a case study of 851 cases, mucormycosis had a reported mortality of 46% (Jeong et al., 2019). Despite advancements in diagnosis and treatment methods, a rise in incidence of mucormycosis is of increasing concern as underlying risk factors fuel pathogenesis. *M. hiemalis* is not a typical cause of mucormycosis infection due to its low temperature tolerance of 33°C, but at least five cases have been diagnosed to date (Lu et al., 2013; Prabhu & Patel, 2004). Three cases of cutaneous *M. hiemalis* infections have been diagnosed: two instances of *M. hiemalis* infection were identified in young immunocompetent girls, and one in a 76-year-old woman (Jin et al., 2015; Paes De Oliveira-Neto et al., 2006; Prevoo et al., 1991). Subcutaneous infections by *M. hiemalis* have been identified in a 44-year-old immunocompetent patient and an immunodeficient diabetic patient (Costa et al., 1990; Desai et al., 2013). More recently, an increase in *Aspergillus* and *Mucor* infections has been observed in patients with COVID-19, showing a coinfection dynamic between fungi and virus (Marr et al., 2021; Pal et al., 2021). The increase in risk and incidence of fungal diseases like mucormycosis highlights the need for the development of antifungal therapies and understanding of immune mechanisms through research on fungal-host interactions.

We present novel research establishing *C. briggsae* as a natural host of the fungal pathogen *M. hiemalis.* Our results demonstrate that *M. hiemalis* is ingested by *Caenorhabditis* nematodes, where it colonizes the intestine with vegetative growth and causes intestinal perforation and eventual death. We show that *M. hiemalis* infects *Caenorhabditis* nematodes with a wide host range that includes *C. briggsae, C. elegans,* and several more distantly related species. We characterized the pathogenesis of *M. hiemalis*, showing that the fungal spores are ingested and then germinate into hyphae and chlamydospores within the nematode intestine causing physical damage to the intestinal barrier. Finally, we assessed gene expression changes for a selection of genes associated with innate immunity mechanisms in *C. elegans* and found that *M. hiemalis* does not induce these previously identified immune response genes. Our findings characterize a new naturally occurring host-pathogen relationship between *Caenorhabditis* nematodes and *M. hiemalis*, which opens new avenues for studying host immune responses and pathogenesis factors of Mucorales fungi using nematode models.

## Results

### Novel intestinal fungus *Mucor hiemalis* identified in wild *Caenorhabditis briggsae*

We first isolated *Mucor hiemalis* and its host *Caenorhabditis briggsae* from a leaf litter sample from Ridgway, Pennsylvania, USA on September 30, 2023. Co-culturing the wild nematodes with IPR reporter *C. elegans* resulted in *pals-5p*::GFP expression, which typically indicates infection transmission (Figure 1A). We observed nematodes grown on the fungus showed a distended intestinal phenotype after staining with the chitin binding dye Direct Yellow 96 (DY96). Microscopy revealed that the fungus is ingested by nematodes. Vegetative growth of the fungus was observed inside of the intestinal lumen of *C. briggsae,* and some fungus colonized the body cavity of dead nematodes. As *C. elegans* infection by other dimorphic and filamentous fungi including *Cryptococcus neoformans* and *Candida albicans* were previously studied, we pursued sequencing methods to identify the pathogen. Sanger sequencing and BLASTN search of the internal transcribed spacer 1 (ITS) gene of the wild isolate JNSP1 revealed a high sequence similarity with other *M. hiemalis* isolates. Other ITS sequences of *Mucor* isolates were obtained, and the resulting phylogenetic tree revealed a distinct clade of *M. hiemalis* including the JNSP1 isolate from Nematode Hunters (Figure 1B). The sequencing of the wild host nematode cultured with *M. hiemalis* was determined to be *Caenorhabditis briggsae* by small-subunit ribosomal RNA and internal transcribed spacer 2 sequencing.

**Figure 1:**
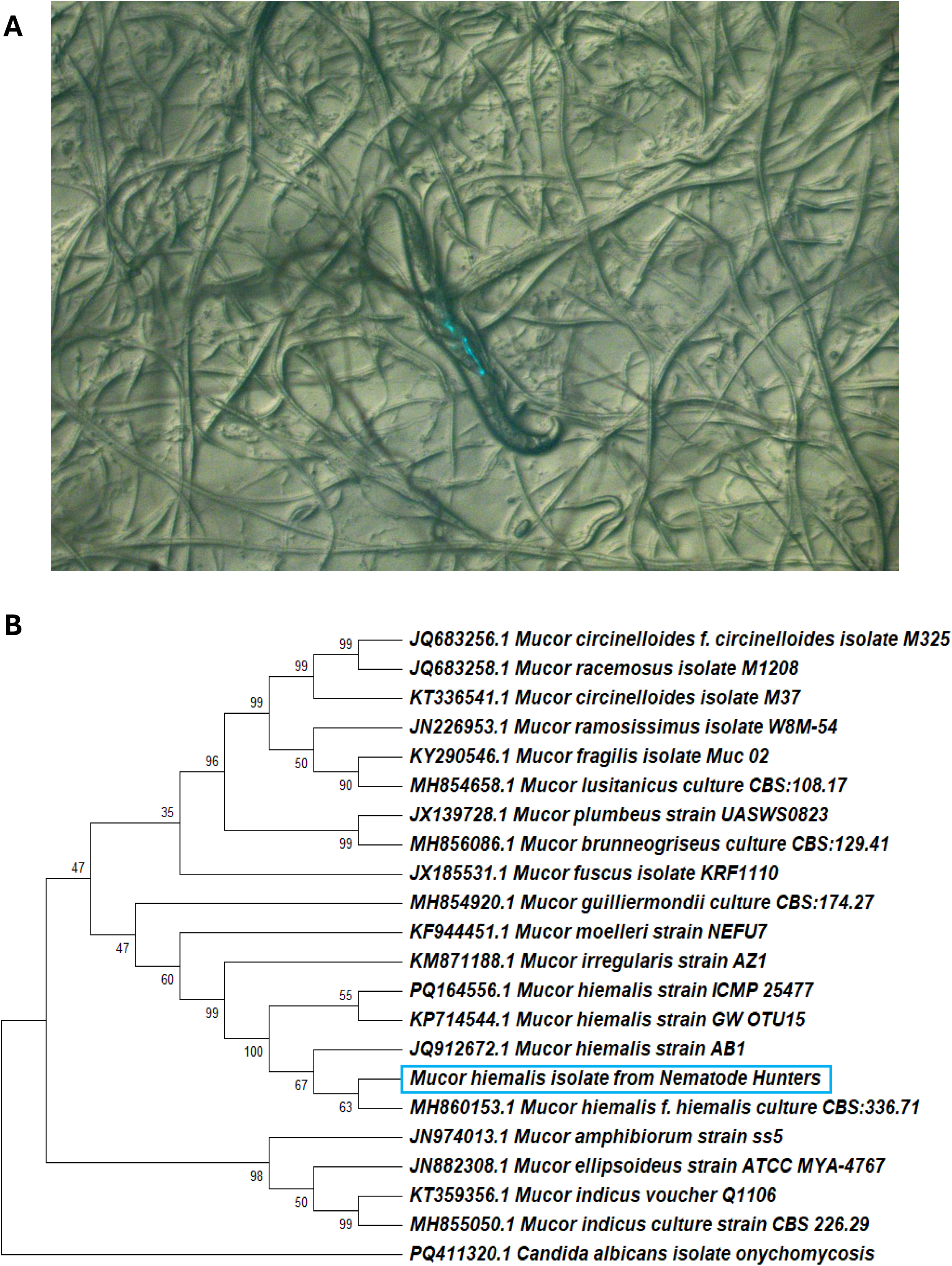
Identification of *Mucor hiemalis* and *Caenorhabditis briggsae* by ITS sequencing. (A) Co-culture experiment performed with *C. briggsae* SOW21 wild isolate and SOW6 *pals-5p*::*GFP* IPR infection reporter *C. elegans* showing positive *pals-5::GFP* expression. (B) Evolutionary relationship of *Mucor* sp. in relation and similarity to *M. hiemalis* isolated from leaf litter sample. Mucor sequences were gathered from NCBI BLASTN. Phylogeny inferred using maximum likelihood method and Hasegawa-Kishino-Yano (1985) model of nucleotide substitutions and the tree with the highest log likelihood (−4,797.06) is shown. The evolutionary rate differences among sites were modeled using a discrete Gamma distribution across 4 categories (*+G*, parameter = 0.5606). The analytical procedure encompassed 22 coding nucleotide sequences using 1st, 2nd, 3rd, and non-coding positions with 746 positions in the final dataset. Evolutionary analyses were conducted in MEGA12.

### Ingested *M. hiemalis* spores germinate hyphal bodies and accumulate inside of the *Caenorhabditis* nematode intestine

Next, we characterized the life cycle of *M. hiemalis* when ingested by nematode hosts with staining and microscopy methods (Figure 2). Reproductive stages of *M. hiemalis* were identified within nematode hosts including spores (S), chlamydospores (C), and filamentous hyphae (H) (Figure 2A-2D). The infection of *Caenorhabditis* hosts begins with spore ingestion as spores fill the intestinal lumen (Figure 2A). Spores germinate into hyphae and chlamydospores and accumulate in the posterior end of the nematode (Figures 2B & 2C). Later time points of fungal accumulation revealed fungus germinating outside of the intestinal lumen and throughout the body cavity of the nematode (Figure 2D). Fungal stages were characterized across five-day time points for a range of nematode growth stages (Figures 2E-2G). When *M. hiemalis* was introduced to larval stage 4 (Figure 2E) *C. elegan*s, spores were first observed in the intestine on the third day after the introduction of spores. In day 1 adult nematodes, spores were observed on the first day (Figure 2F). We attributed the difference in spore ingestion between the L4 nematodes and day 1 adults to nematode age and size differences. *C. elegans* mouth size is about 4 µm at L4 stage and it grows larger with nematode age (Gubert et al., 2023). We measured *M. hiemalis* spores to be ~6 µm in diameter and the L4 nematodes had grown to a larger adult stage by the third day of observation. Spore counts decreased after initial ingestion in all stages as spores germinated into alternative growth forms. A high frequency of nematodes contained hyphae and chlamydospores by the fourth and fifth days of infection (Figures 2E-2G, Figure S1). A difference in *M. hiemalis* ingestion was observed in adult day 4 nematodes, as no spores were observed in the intestine until the third day (Figure 2G). We attributed the reduction of ingestion to reduced pharyngeal pumping due to age-related muscle degeneration and physical accumulation of fungus inside of the pharynx (Huang et al., 2004).

**Figure 2:**
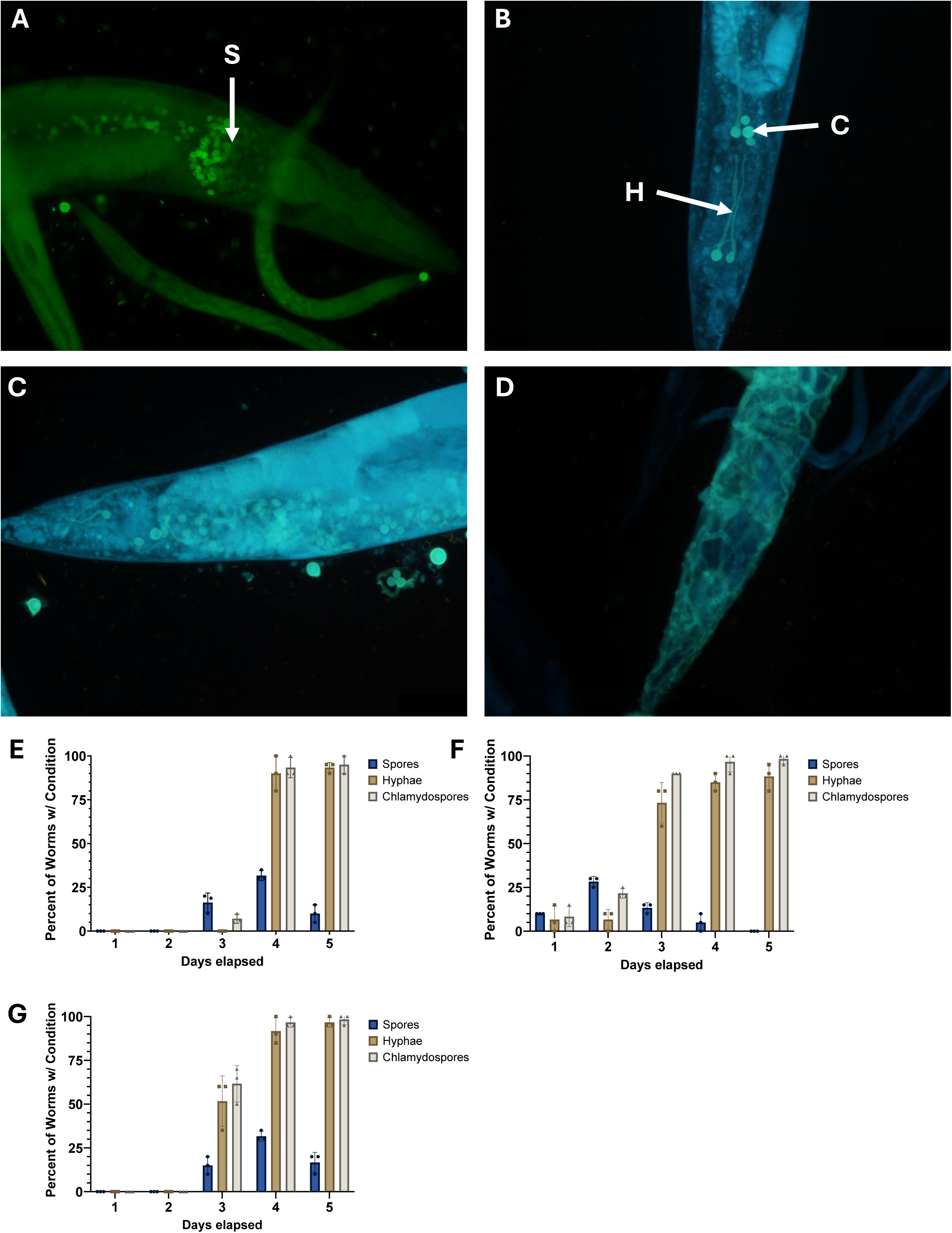
Ingested *M. hiemalis* spores germinate hyphal bodies and accumulate inside of the *Caenorhabditis* nematode intestine. Fluorescence micrographs of *M. hiemalis* growth stages stained with Direct Yellow 96 in infected wild type N2 *C. elegans*. Nematodes were scored for fungal growth conditions in biological triplicates. Percentage of *C. elegans* containing *M. hiemalis* spores (S), hyphae (H), and chlamydospores (C) were quantified. (A) *M. hiemalis* spores are ingested by *Caenorhabditis* nematodes and accumulate in the intestinal lumen. (B) Vegetative growth from spores to filamentous hyphae at the posterior end of the intestine. Chlamydospores arise from hyphal segments during asexual spore production. (C) Accumulation of vegetative growth forms in the posterior end of the nematode. (D) Hyphal growth breaching the intestinal lumen and into the nematode body cavity. (E) *M. hiemalis* growth stage counts in N2 *C. elegans* across 5 days of observation by fluorescence microscopy with (E) larval stage 4, (F) adult day 1, (G) adult day 4 nematodes. Graphs show the average percentage of nematodes showing each fungal stage from 3 independent experiments combined (n = 20 for each experiment). Error bars show standard deviation.

### Caenorhabditis nematodes prefer E. coli mixed with M. hiemalis spores

Given that we observed nematodes ingesting the fungus, we investigated the preference of *Caenorhabditis* nematodes when presented with *M. hiemalis* or alternative food choices. Nematodes can exhibit preference toward high quality food that supports growth and avoidance of foods that do not support growth (Shtonda & Avery, 2006). Nematodes have also demonstrated pathogen avoidance after initial ingestion by olfactory learning with *P. aeruginosa* (Zhang et al., 2005) and have demonstrated lawn-leaving behavior with the pathogenic bacteria *Serratia marcescens* by chemosensation (Pradel et al., 2007). We assayed if *C. elegans* displayed behavioral preference toward *M. hiemalis* through a choice assay (Figure 3A). Preference indexes were calculated and analyzed with a one-sample t-test to a theoretical mean of 0, where 0 indicates no preference. Nematodes did not show a significant preference toward OP50-1 *E. coli* compared to *M. hiemalis* spores alone (Figure 3B). *C. elegans* showed preference to the mixture of *M. hiemalis* and OP50-1 compared to *M. hiemalis* or OP50-1 alone (p = 0.0102, one-sample t-test). Interestingly, the mixture of fungal and bacterial components is more comparable to nematode food sources in natural ecosystems than isolated bacterial strains. The spore mixture with OP50-1 was utilized for infection protocols in future experiments.

**Figure 3:**
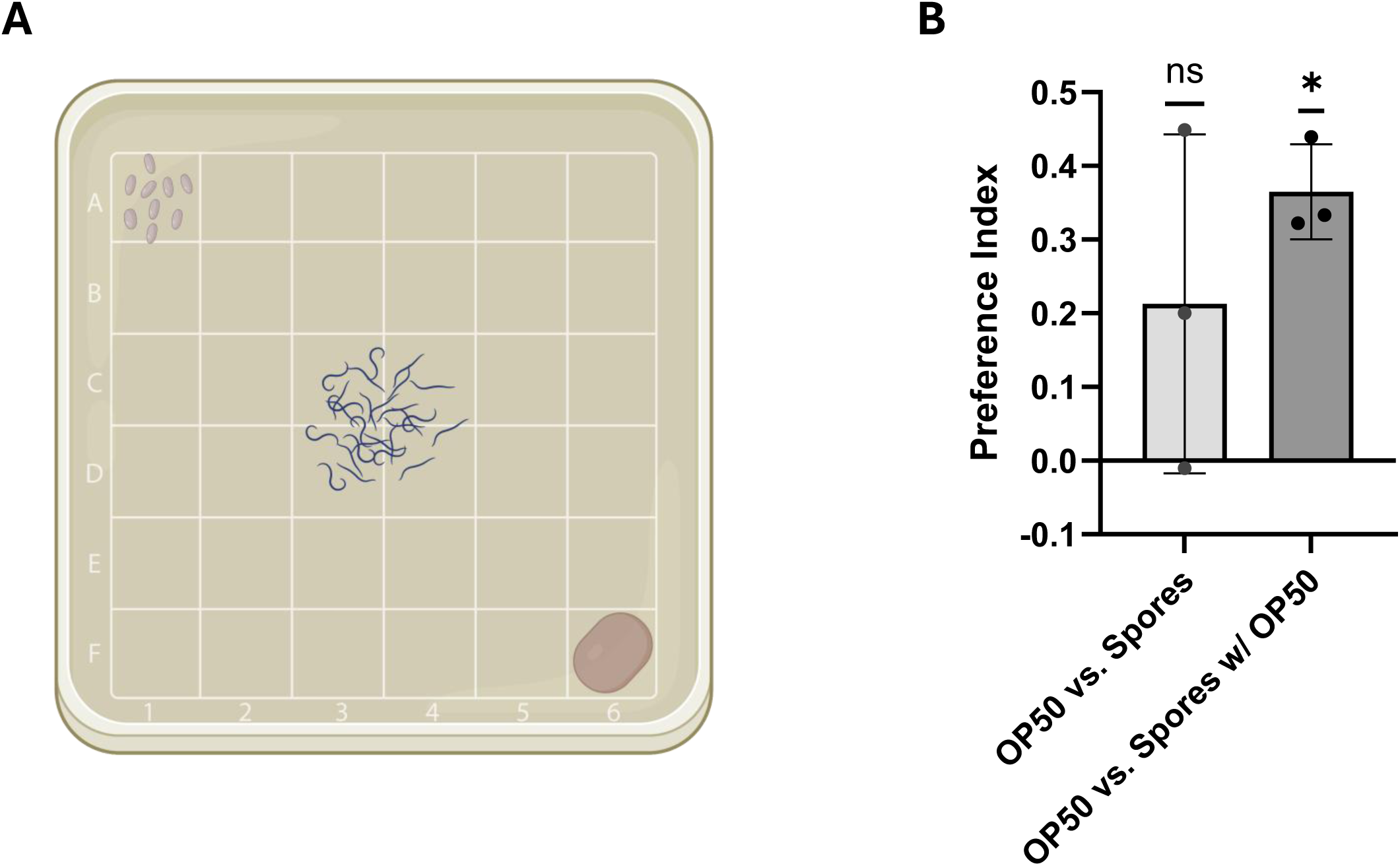
*Caenorhabditis elegans* show preference for ingesting *M. hiemalis* spore and *E. coli* mixture. Preference indexes were calculated from triplicate experiments tested in parallel using 100 nematodes per group. A one-sample t-test was used to compare the preference score to no preference. (A) A schematic representation of assay plate used for calculating preference indexes. *M. hiemalis* spores were placed in the top left corner and OP50-1 *E. coli* was placed in the bottom right corner. 100 *C. elegans* were introduced in the middle four squares of the plate. (B) Food preference index of *M. hiemalis* spores and spore mixture compared to OP50-1. *C. elegans* showed no preference between fungal spores in water compared to OP50–1 (one-sample t-test, p = 0.2500, n=3). Nematodes showed a significant preference toward a fungal mixture of spores and OP50-1 over OP50-1 alone (one-sample t-test, p = 0.0102, n = 3). Experiment was repeated 3 times independently. Error bars show standard deviation.

### *M. hiemalis* kills *Caenorhabditis* nematodes with a wide host range

After confirming *M. hiemalis* ingestion in nematodes, we assayed the survivability of nematodes to observe if *M. hiemalis* infection reduced their lifespan. We performed lifespan analysis on a range of species including our original wild isolate of *C. briggsae* (Figure 4A), *C. elegans* (Figure 4B), *C. japonica* (Figure 4C), and *C. tropicalis* (Figure 4D). These species are relatively distantly related, sharing a common ancestor more than 80 mya (Baker & Woollard, 2019). *M. hiemalis* significantly reduced the survivability of all species tested with similar percent reductions in median survival (Figure 4E, Figure S2). *C. briggsae* on infection plates lived to a median survival of 8.50 days compared to 12 days on control OP50 plates. *C. elegans* on infection plates lived to a median survival of 7.50 days compared to 11.50 days on control OP50 plates. *C. japonica* on infection plates lived to a median survival of 8.00 days compared to 12.00 days on control OP50 plates. *C. tropicalis* on infection plates lived to a median survival of 7.00 days compared to 11.00 days on control OP50 plates. *M. hiemalis* reduced the survivability of all species tested with similar percent reductions in median survival (*C. briggsae* 29.2%; *C. elegans* 34.8%; *C. japonica* 33.3%; *C. tropicalis* 36.4%). A comparison of survival curves using a Log-rank (Mantel-Cox) test revealed statistically significant reduction in all species compared to controls (p < 0.0001). *M. hiemalis* consistently reduced the lifespan of all four species of nematodes, suggesting that it has a wide host range and consistent pathogenesis.

**Figure 4:**
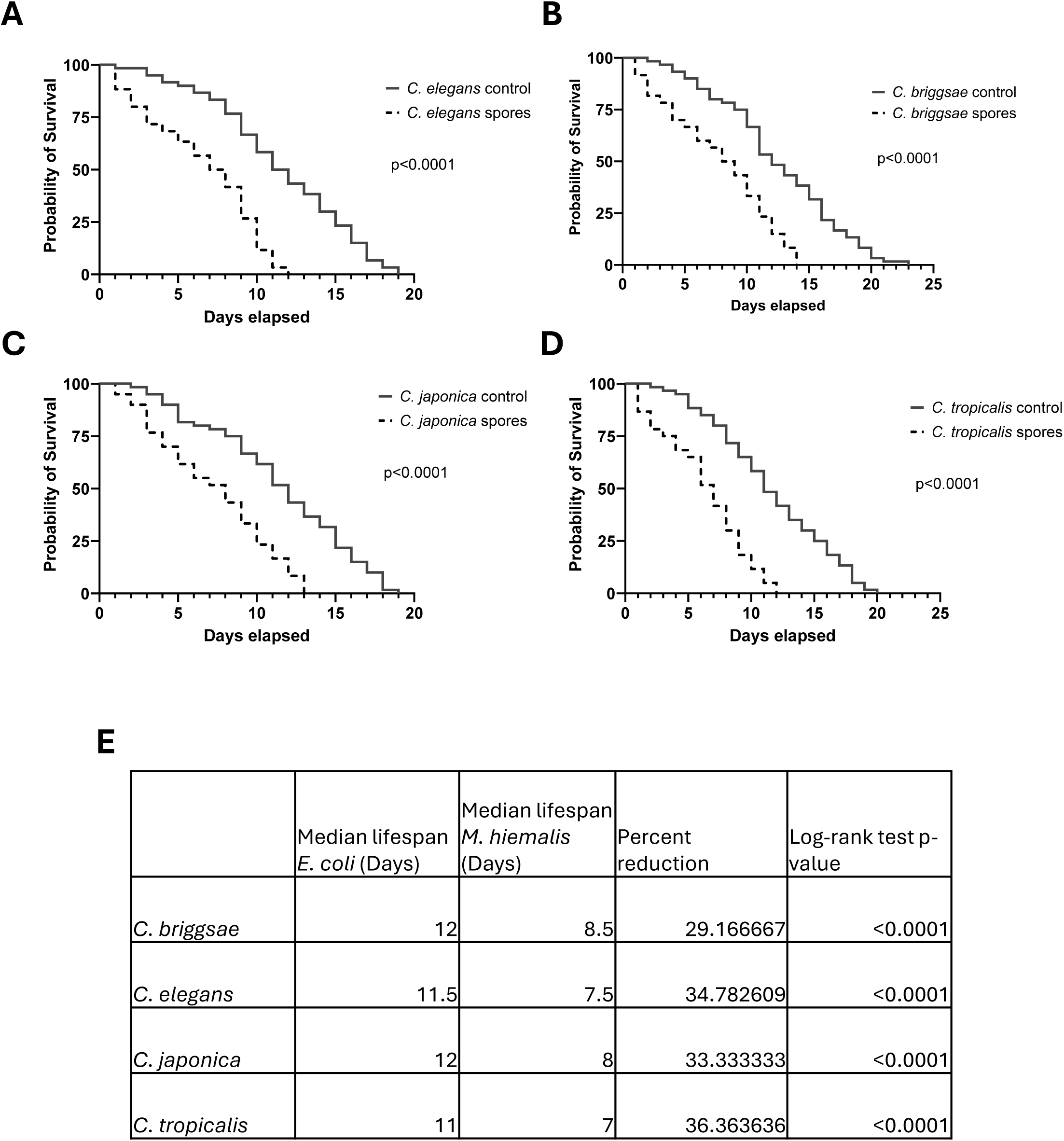
*M. hiemalis* kills *Caenorhabditis* nematodes with a wide host range. Survival assays of (A) *C. elegans* N2, (B) *C. japonica* DF5081, (C) *C. tropicalis* JU1373, and (D) *C. briggsae* SOW21 wild isolate grown on *M. hiemalis* + OP50 infection plates or OP50 control plates. Infection by *M. hiemalis* significantly reduced lifespan in each species tested (log-rank test, p < 0.001). Graphs show combined lifespan from three independent biological replicates (n = 20 for each experiment). (E) Table showing median lifespan and percent reduction with Log-rank (Mantel-Cox) tests comparing lifespans of nematodes grown with vs. without *M. hiemalis*.

### *M. hiemalis* infection kills nematodes by perforating the intestinal barrier

To better understand the mechanism by which *M. hiemalis* kills nematodes, we examined the intestinal barrier integrity relative to fungal growth. The previous microscopy experiment revealed *M. hiemalis* growth outside of the intestinal lumen at later time points, so we hypothesized that *M. hiemalis* growth perforates the intestine. To examine the intestinal barrier integrity, we fed nematodes with propidium iodide (PI) stain. This stain remains contained within the intestinal lumen of intact intestines but will leak into the body cavity in nematodes with a perforated intestine (Figure 5A). Nematodes were scored positive for intestinal perforation if the body cavity was stained (Figure 5B). No PI staining occurred on the first day in either control or infection plates. On the second day, 5% of control nematodes showed body cavity staining by PI compared to 22% of infected nematodes. PI staining occurred in control nematodes likely due to cellular degeneration associated with increasing age. Nematodes infected with *M. hiemalis* had significantly more nematodes with PI staining of the cavity compared to controls on the third (p = 0.0002, unpaired t-test) and fourth day (p = 0.0002, unpaired t-test). As we observed PI staining in infected nematodes, we concluded that accumulation of *M. hiemalis* in the intestine induces perforation.

**Figure 5:**
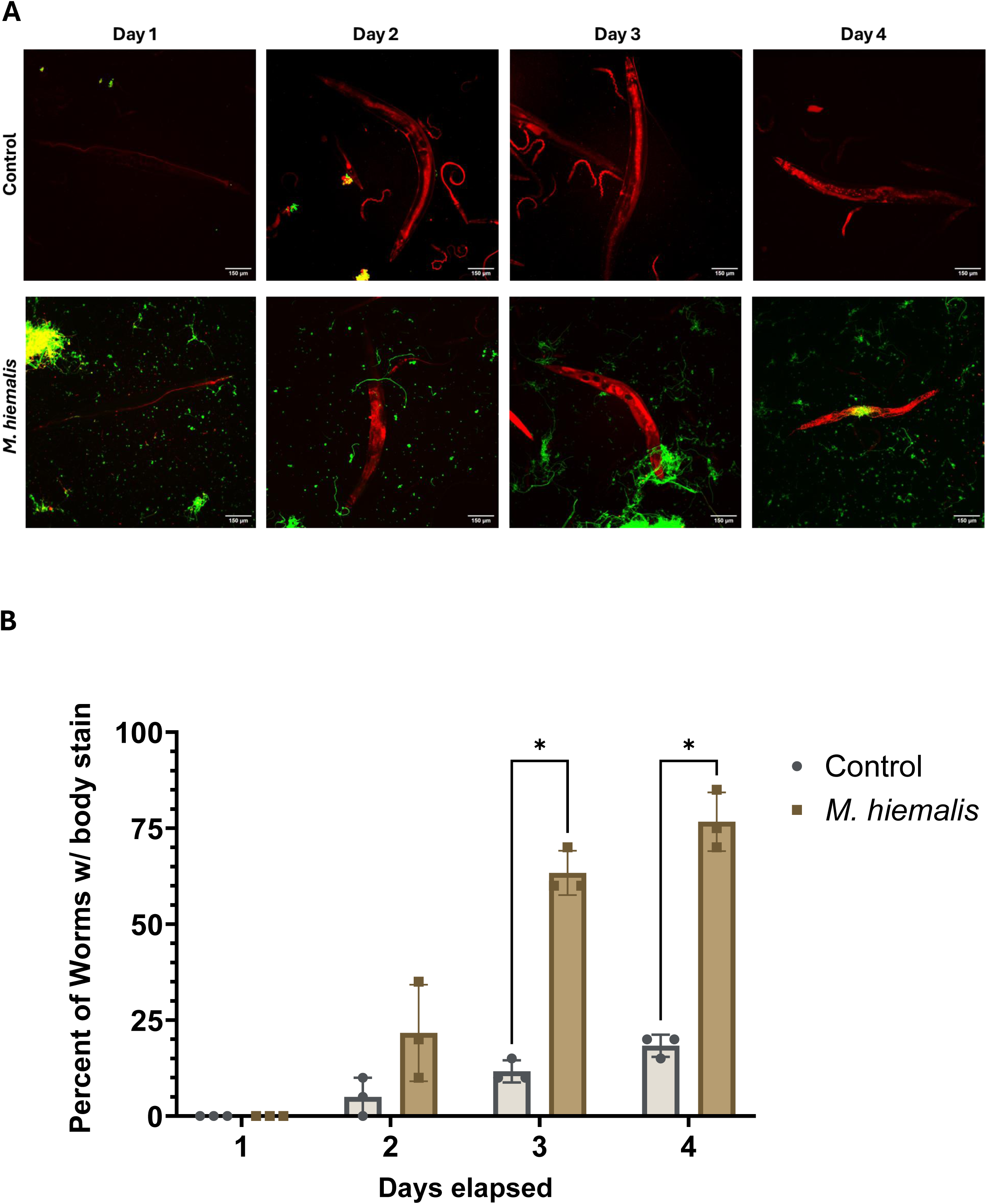
*M. hiemalis* infection perforates the intestinal barrier. (A) Fluorescence micrographs of N2 *C. elegans* grown on *M. hiemalis* infection plates and OP50 control plates after DY96 and PI staining. PI staining outside of the intestinal lumen indicates that the intestinal barrier is no longer intact. (B) Percentages of infected N2 nematodes and controls with PI staining in the body cavity. Nematodes were scored positive for PI staining if fungus was present and PI staining occurred outside of the intestine. Graph is the average of 3 independent experiments (n = 20). Error bars show standard deviation.

### IPR and other immunity genes are not upregulated in response to *M. hiemalis* infection

Next, we investigated common nematode immune responses to determine if any were induced by *M. hiemalis* infection. Since *C. elegans* lack adaptive immunity, these nematodes rely on innate immune mechanisms to protect and clear infections (Tran & Luallen, 2024). The *C. elegans* intracellular pathogen response (IPR) plays an important role in providing intestinal immunity against Orsay virus and the microsporidia *N. parisii* (Reddy et al., 2017, 2019). *C. elegans* also have an established p38 MAP kinase pathway that contributes to innate immunity (Kim et al., 2002). In response to some fungal pathogens, *C. elegans* produces antimicrobial peptides through a TGF-β signaling pathway independent of the p38 pathway (Zugasti & Ewbank, 2009). Lastly *C. elegans* can respond to pathogen infection by inducing the ZIP-2/IRG-1 pathway to respond to toxins (Dunbar et al., 2012). We assessed activation of common *C. elegans* immune responses during *M. hiemalis* infection using a collection of IPR genes and other innate immunity genes associated with p38 MAPK, TGF-β, and zip-2/irg-1 pathways (Figure 6). qRT-PCR gene expression analysis showed the majority of the IPR and innate immunity genes in larval stage 4 and adult day 2 *C. elegans* were not significantly induced by *M. hiemalis* infection. A one-sample t-test was performed to compare gene expression fold change to a theoretical mean of 1, indicating no change. In larval stage 4 nematodes, the gene expression of *irg-1* was significantly decreased (p = 0.0305). The reduction of *irg-1* was unexpected as previous studies found that *irg-1* is a transcriptional response to the bacterial pathogen *P. aeruginosa*. The cause for the reduction of *irg-1* in young nematodes is unknown. In adult day 2 nematodes, *pals-5* showed a nearly significant increase (p = 0.0518) and *eol-1* showed a nearly significant reduction (p = 0.0514). The other innate immunity genes were not significantly changed. Multiple unpaired t-tests analyzed the differences in gene expression fold change between larval stage 4 nematodes and adult D2 nematodes. No significant changes in gene expression were found between the two nematode growth stages. Although none of the genes we examined were upregulated in response to *M. hiemalis,* further investigation could provide insight into novel immunity responses to fungal pathogens.

**Figure 6:**
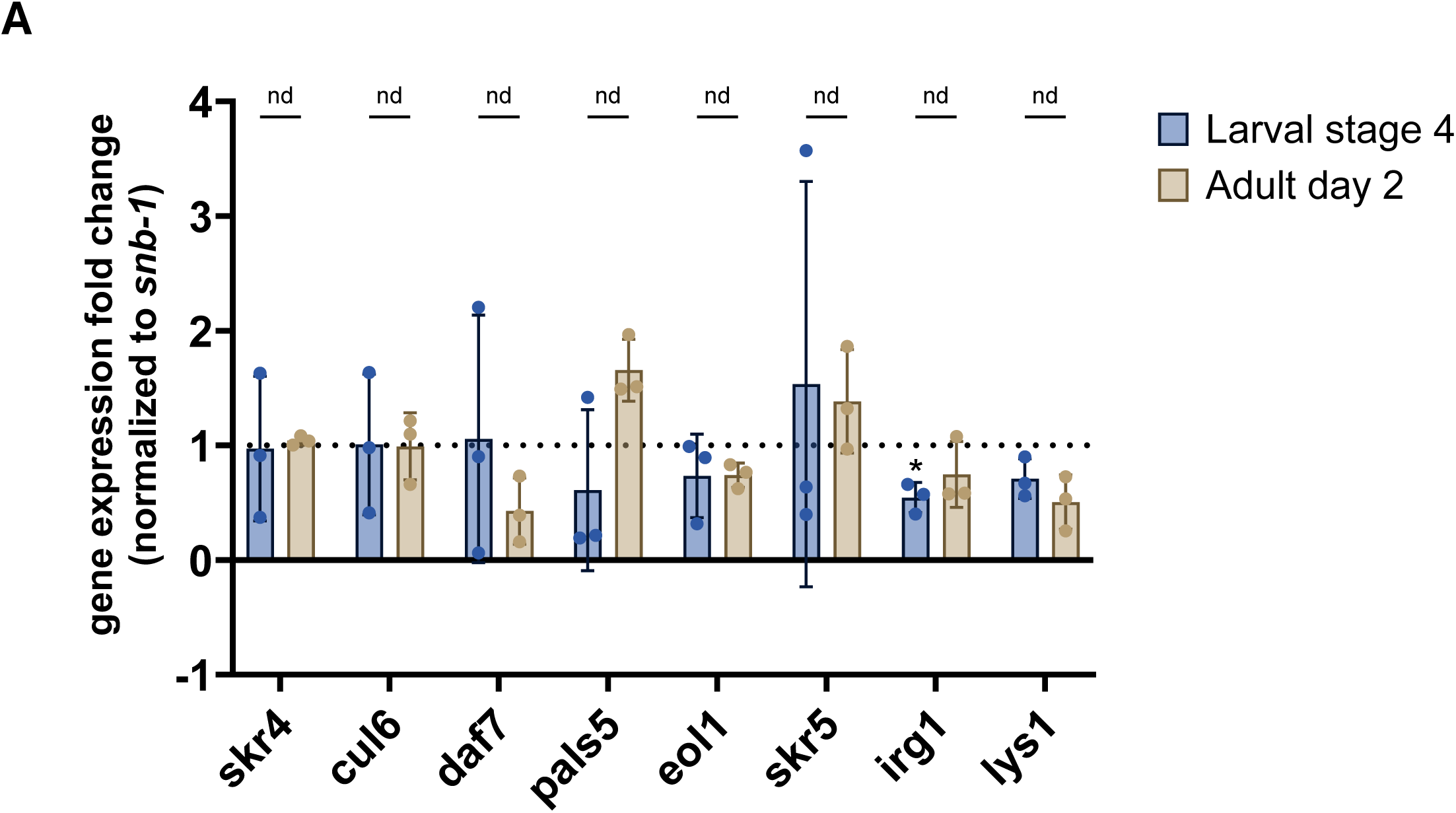
IPR and other immunity genes are not upregulated in response to *M. hiemalis* infection. 1000 synchronized larval stage 4 and adult day 2 N2 *C. elegans* were incubated on fungal infection plates and OP50-1 control plates for 2 days before RNA extraction. Experiment was repeated 3 times independently. Relative fold changes in gene expression were normalized to *snb-1* and compared to control groups. Graph shows gene expression comparison between larval stage 4 and adult day 2 nematodes infected with *M. hiemalis*. Error bars show standard deviation. Significance was determined by multiple unpaired t-tests comparing means of larval and adult nematodes. Individual gene expression values were compared to a value of 1 with one-sample t-tests. No significant differences were observed in gene expression between the two nematode growth stages. IPR genes *skr-4, cul-6, pals-5,* and *eol-1* are not induced during *M. hiemalis* infection in larval stage 4 or adult day 2 *C. elegans*. Innate immunity genes *daf-7, skr-5* and *lys-1* are also not induced. *irg-1* gene expression is reduced upon *M. hiemalis* infection in larval stage 4 nematodes (p = 0.0264, n = 3).

## Discussion

The study of host-pathogen interactions using the *C. elegans* model is limited by the number and variety of naturally occurring nematode pathogens that have been identified. Here we identified the human-pathogenic dimorphic fungus *M. hiemalis* as a natural fungal pathogen of *Caenorhabditis* nematodes and developed a system to model fungal pathogenicity. We characterized the life cycle of *M. hiemalis* in nematode hosts and found that infection begins with spore ingestion and vegetative growth accumulates in the posterior end of the nematode. Nematodes showed preference toward a mixture of *M. hiemalis* spores and OP50-1. We observed a significant reduction in survivability in multiple *Caenorhabditis* species and determined that *M. hiemalis* kills nematodes by perforating the intestine. We also did not observe any significant induction of the common nematode immune responses in response to the fungal infection. In total, we have characterized a novel fungal pathogen of *Caenorhabditis* nematodes and provide insights into their host-pathogen relationship.

To investigate the feeding preferences of *Caenorhabditis* hosts, we designed an assay to determine whether they preferred *M. hiemalis* or a standard laboratory food source, OP50-1 *E. coli*. Our observations revealed a significant preference for a mixture of *M. hiemalis* spores and OP50-1 compared to OP50-1 alone, which we subsequently used in further experiments. Interestingly, nematodes did not show a significant preference for *M. hiemalis* alone, yet in both conditions involving the fungus, nematodes showed a positive average preference. *C. elegans* employs chemosensing to detect chemical signals and secondary metabolites from surrounding microbes, and they can distinguish between high and poor quality food (Bargmann, 2006; Shtonda & Avery, 2006). Our findings raise questions about whether *C. elegans* are detecting metabolites secreted by *M. hiemalis* and whether *C. elegans* sense that *M. hiemalis* can support nematode growth. While we do not know if *C. elegans* derive nutritional value from ingesting *M. hiemalis* spores and fungal structures, it is possible that they are attracted to fungal nutrients or fermentation byproducts. Furthermore, *M. hiemalis* might secrete chemicals that mimic other processes that are attractive to *C. elegans*. For instance, the nematode-trapping fungus *Arthrobotrys oligospora* attracts nematodes by secreting compounds that mimic sex and food cues (Hsueh et al., 2017). Olfactory mimicry could potentially play a role in the initial attraction of *C. elegans* to *M. hiemalis*.

Given that specific behaviors are linked to the initial attraction of *C. elegans* to food sources, it is plausible that these nematodes could also develop learned avoidance towards *M. hiemalis* following initial ingestion. *C. elegans* have been shown to avoid certain pathogenic bacteria such as *S. marcescens* and *P. aeruginosa* after an initial encounter. When fed *S. marcescens*, *C. elegans* recognizes serrawetin, a chemical secreted by the bacterium and subsequent detection by G protein and toll-like pathways are responsible for lawn-leaving behavior (Pradel et al., 2007). A similar interaction is observed with *P. aeruginosa,* where *C. elegans* exhibits transgenerational avoidance behaviors by recognizing bacterial small RNAs (Sengupta et al., 2024). Our current study only captured the initial attraction of *M. hiemalis* due to the use of sodium azide to immobilize the nematodes after their first choice. Future investigations could reveal whether nematodes also develop learned avoidance behaviors towards *M. hiemalis*.

Our analysis did not reveal a significant upregulation of common innate immunity genes typically associated with viral, bacterial, and fungal pathogens of *C. elegans*. Although we observed a near significant upregulation of *pals-5* (p = 0.0518) in adult day 2 nematodes, the level of *pals-5* expression detected by qPCR was lower than the anticipated levels based on our co-culture assays. Several factors could account for this discrepancy. In co-culture tests, *pals-5p::GFP* expression typically became apparent after three days post infection. The qPCR experiment to quantify gene expression was conducted after only two days of infection. Extending the qPCR analysis to three or four days post-infection might reveal higher *pals-5* expression levels. Additionally, the presence of excess progeny prior to RNA extraction could have diluted the expression of innate immunity genes as normalization with *snb-1* was performed relative to the total amount of DNA per extraction. IPR genes are categorized based on their dependence on the transcription factor ZIP-1 and multiple pathways can activate the IPR to induce *pals-5p::GFP* (Lažetić et al., 2022). While intracellular infection by Orsay virus and microsporidia, proteasome inhibition, or prolonged heat stress are known inducers of the *C. elegans* IPR, it is possible that *M. hiemalis* could activate *pals-5* through a distinct pathway (Lažetić et al., 2022). Future studies could investigate the ZIP-1 dependence of *pals-5::GFP* activation by *M. hiemalis*, as well as a broader survey of IPR gene expression.

We did not observe activation of previously characterized *C. elegans* innate immune pathways associated with fungal infections, including the insulin/insulin-like growth factor 1, DAF-16, and p38-MAPK signaling cascades. The absence of these responses is significant, suggesting that *M. hiemalis* might elicit a unique host defense mechanism in *C. elegans*. In contrast, humans mount both innate and adaptive immune responses to invasive Mucorales infections. These fungi often establish infection by evading the initial defenses of host epithelial tissues by entry through traumatic wounds or via ingestion (Skiada et al., 2012; Spellberg, 2012). The specific growth and developmental stage of Mucorales also influences the host immune effectors involved. For instance, macrophages and neutrophils detect Mucorales growth following spore detection in the host epithelium (Ghuman & Voelz, 2017). Macrophages can inhibit spore germination into hyphae and other growth structures, but they are unable to eliminate resting spores (Ghuman & Voelz, 2017). Natural killer cells also play a regulatory role in the human immune response to Mucorales. Cytokines like IL-12 and type-1 interferons activate natural killer cells and they co-localize with macrophages at infection sites and areas of tissue damage (Ghuman & Voelz, 2017). These innate immune effectors play a crucial role in initiating the adaptive immune response in humans where T cells provide defense against the fungal pathogen through the secretion of antifungal cytokines and defensins (Ghuman & Voelz, 2017). *C. elegans* lacks an adaptive immune system and circulating immune cells such as macrophages and neutrophils, so its detection of *M. hiemalis* may rely on unique PAMP or DAMP-like receptors capable of recognizing toxins or damage associated with *M. hiemalis* infection. Notably, the *C. elegans* IPR shares similarities with mammalian type-I interferon responses (Lažetić et al., 2023). Although the mechanism is unclear, components of the IPR might secrete signaling proteins to activate a two-step response to infection similar to the type-1 interferon pathway (Lažetić et al., 2023). These parallels between conserved pathways present an exciting opportunity to characterize a novel *C. elegans* immune response to *M. hiemalis*.

## Conclusion

In conclusion, this study identified the human-pathogenic dimorphic fungus *M. hiemalis* as a novel, naturally occurring pathogen of *Caenorhabditis* nematodes, establishing *Caenorhabditis* as a model for dissecting this host-pathogen interaction. We characterized the fungal life cycle inside of nematode hosts, assayed the reduction of nematode lifespan across diverse species, and investigated the host’s potential immune responses to infection. Our findings demonstrate that a wide range of species ingest *M. hiemalis* spores and subsequent fungal growth kills nematodes by perforating the intestinal barrier. Our investigation of known *C. elegans* immune pathways revealed a surprising absence of immune gene upregulation, suggesting the possibility of uncharacterized defense mechanisms employed by the nematode against fungal infection. This research offers novel insights into the interactions between *Caenorhabditis* nematodes and *M.* hiemalis, which expands the collection of natural nematode pathogens and opens new avenues for research of host-fungus interactions.

## Acknowledgements

We thank Erik Andersen and the CaeNDR project for their sequencing support of wild *Caenorhabditis* strains. We thank Emily Troemel for sharing *C. elegans* strains. Some strains were provided by the CGC, which is funded by NIH Office of Research Infrastructure Programs (P40 OD010440). We are grateful to Marcia Raubenstrauch and the students at Francis S. Grandinetti Elementary School for their role in collecting wild nematodes for the Nematode Hunters project. This study was supported by the National Science Foundation under the grant BRC-BIO 2218079.

## Methods

### Co-culture infection transmission experiment

Wild isolate SOW21 *C. briggsae* and SOW6 *C. elegans* were both chunked onto one Nematode Growth Media (NGM) plate with a flame sterilized spatula. Co-culture plates were maintained at 20°C and checked daily for reporter fluorescence compared to SOW6 only controls. Positive co-culture tests were imaged on an Olympus SZX10 stereo microscope at 6X magnification. Images were captured with an Olympus DP23 camera on Olympus cellSENS imaging software (https://evidentscientific.com/en/).

### Sequencing of Mucor hiemalis and Caenorhabditis briggsae

*Mucor hiemalis* DNA was isolated with a Quick-DNA Miniprep Kit (Zymo Research) according to manufacturer’s instructions. PCR amplification of the internal transcribed spacer region was performed with primers described by White et al., (1990) (ITS1 – 5’-TCCGTAGGTGAACCTGCGG, ITS4 5’-TCCTCCGCTTATTGATATGC). Sanger sequencing was performed by Azenta (South Plainfield, NJ). The resulting sequence of the *M. hiemalis* JNSP1 isolate was submitted to NCBI (GenBank: PV719612). ITS sequences of other Mucor species were obtained from NCBI BLASTN. The phylogeny was inferred using the Maximum likelihood method and Hasegawa-Kishino-Yano model of nucleotide substitutions. The evolutionary rate differences among sites were modeled using a discrete Gamma distribution across 4 categories (*+G*, parameter = 0.5606). The analytical procedure encompassed 22 nucleotide sequences with 746 positions in the final dataset. Evolutionary analyses were conducted in MEGA12 (https://www.megasoftware.net/).

*Caenorhabditis briggsae* (SOW21) DNA was extracted with Quick-DNA Miniprep Kit (Zymo Research). PCR amplification of the 18S ribosomal gene was performed with primers described by Nakacwa et al., (2013) (SSU18A – AAAGATTAAGCCATGCATG, SSU26R – CATTCTTGGCAAATGCTTTCG). Sanger sequencing was performed by Azenta (South Plainfield, NJ). Results were further confirmed with ITS2 sequencing by the CaeNDR project (https://caendr.org/).

### Maintenance of *Caenorhabditis* nematodes

*Caenorhabditis* nematodes were grown and maintained on NGM plates and incubated at 20°C. NGM plates were seeded with *Escherichia coli* OP50-1 bacteria.

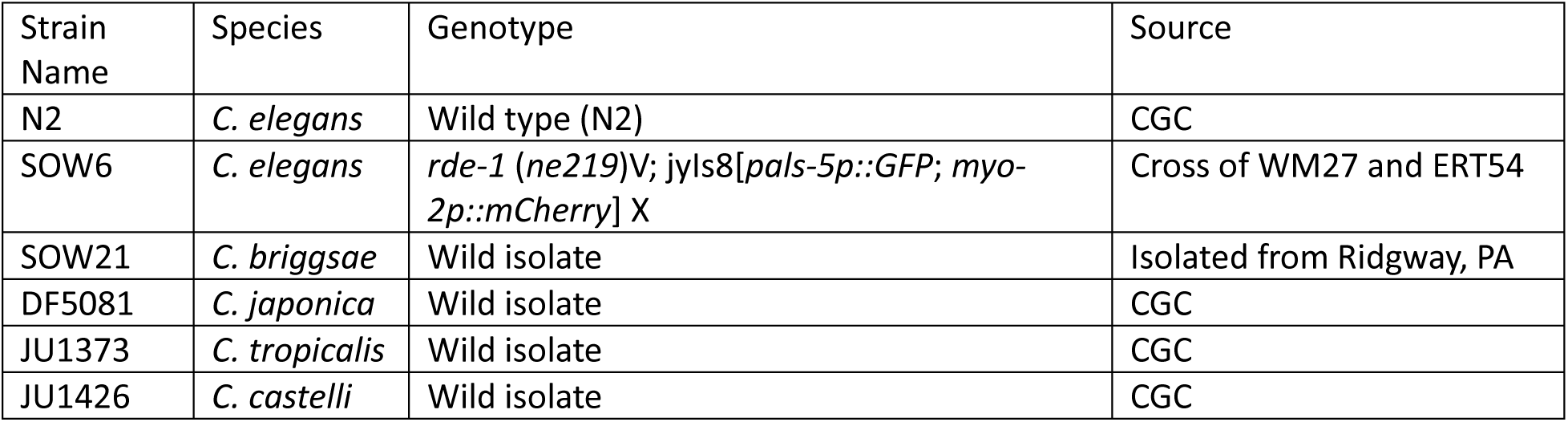

| Strain Name | Species | Genotype | Source |
| --- | --- | --- | --- |
| N2 | *C. elegans* | Wild type (N2) | CGC |
| SOW6 | *C. elegans* | *rde-1 (ne219)V*; *jyls8[pals-5p::GFP; myo-2p::mCherry]* X | Cross of WM27 and ERT54 |
| SOW21 | *C. briggsae* | Wild isolate | Isolated from Ridgway, PA |
| DF5081 | *C. japonica* | Wild isolate | CGC |
| JU1373 | *C. tropicalis* | Wild isolate | CGC |
| JU1426 | *C. castelli* | Wild isolate | CGC |

### Maintenance of Mucor hiemalis

*Mucor hiemalis* fungus were grown and maintained on Yeast Peptone Glucose (YPG) plates. YPG plates were inoculated by transferring spores with a sterile pipette tip in an “X” pattern. Plates were incubated at room temperature with the presence of light to encourage sporangiospore production.

### Spore harvesting and nematode infection

YPG plates were inoculated with *Mucor hiemalis* and grown for four days before spore harvesting. 10mL sterile ddH2O was pipetted onto YPG plates. A sterile cell spreader was used to scrape the top of the mycelial mat to release spores. Plates were tilted to one side and a pipette was used to transfer the spore suspension to a 15mL conical tube. Spore concentrations were determined with a hemocytometer.

Spore solutions were pelleted by centrifugation at 4,300 x g for 10 min and supernatant was discarded. Spore pellets were resuspended with OP50-1 *E. coli* to a dose of 10 million spores per 500 μL. NGM plates were seeded with the spore and OP50-1 mixture and dried in a laminar flow hood. Synchronized nematodes were transferred to fungal plates after plate drying.

### Direct Yellow 96 staining

Infected nematodes were washed off plates into microcentrifuge tubes with M9 + 0.1% Tween-20. Nematodes were washed 2x with M0 + 0.1% Tween-20 and supernatant was discarded. Nematodes were fixed with 700 μL acetone and were incubated overnight or stored at 4°C. Samples were washed with 1 mL 1X PBS + 0.1% Tween-20. 500uL DY96 solution (20 µg/µL DY96, 0.1% SDS in 1x PBS + 0.1% Tween-20) was added and samples were incubated for 30 minutes at 20°C with rotation. Samples were spun down and the excess DY96 solution was discarded. Vectashield (Vector Laboratories) mounting medium was added to samples before mounting on glass slides.

### Fluorescence microscopy

Microscopy was performed on an Olympus BX61 epifluorescence microscope using 10X and 40X objectives. Images were captured using an Olympus DP21 (Figure 2) or a Tucsen Dhyana 400D (Figure 5) camera. µManager (https://micro-manager.org/) software was used to take Z-stacks of nematodes. ImageJ FIJI (https://fiji.sc/) was used to analyze fluorescence images.

### Propidium iodide and Direct Yellow 96 live staining

100 N2 *C. elegans* were synchronized by bleaching and grown to adult day 1. Standardized *M. hiemalis* spore doses were added to each infection plate and nematodes were incubated at 20°C. Nematodes were collected for analysis at 1-4 days after infection. Nematodes were washed off plates into microcentrifuge tubes with M9 + 0.1% Tween-20 and washed 2X with M9. 500 µL DY96 live staining solution (20 µg/µL DY96, 1xPBS + 0.1% Tween-20) and 30 μL propidium iodide (Thermo Scientific) was added. Samples were incubated for 1 hour at 20°C with rotation. Samples were spun down and the excess dye was discarded. Samples were washed 2X with PBST. Vectashield (Vector Laboratories) mounting medium was added to samples before mounting on slides. Image J FIJI was used to view image stacks. Nematodes were scored positive for intestinal perforation if cells outside of the intestinal lumen were stained with PI. 20 nematodes were scored per trial, and experiment was repeated three times independently.

### Survival assay

Survival assays to determine the host range of *M. hiemalis* were performed with the following species: *C. elegans* (N2), *C. japonica* (DF5081), *C. tropicalis* (JU1373), and *C. briggsae* (SOW21). Survival assays were carried out in biological triplicates with spore treatments and controls. Nematodes were synchronized by bleaching and grown to larval stage 4. 20 nematodes were transferred to either infection plates with spore + OP50-1 mixture or control plates seeded with OP50-1. Nematodes were tested for survival every day by prodding with a platinum wire. Nematodes were transferred to new plates every 2 days to prevent scoring interference by progeny. Statistical analysis of nematode survival was performed with Log-rank (Mantel-Cox) analyses in Graphpad Prism v10 software.

### Food preference assay

Square petri dishes with grids were prepared with NGM media. Preference assay plates were inoculated with OP50-1 and spores/spore mixture with OP50-1 as shown in Figure 3. Spore conditions were at a dose of 1 million spores in either 200 µL sterile ddH2O or OP50-1. Plates were dried in a laminar flow hood and 100 µL sodium azide (0.5M) was added to each choice to prevent travel between choices. About 100 synchronized adult day 1 N2 *C. elegans* were placed in the middle four squares on the choice plate. A Kimwipe was used to remove excess M9 buffer. Nematodes were counted after 1 hour and were scored based on location and choice. Experiments were carried out in biological triplicate. Preference indexes per trial were calculated and averaged. Statistical analysis of nematode choice preference was performed in Prism software. Means were analyzed with a one-sample t-test against a theoretical mean of zero.

### Immune response quantification by qRT-PCR

1000 N2 *C. elegans* were synchronized and grown in biological triplicates. Infection plates were inoculated with spore mixture at either larval 4 or adult day 2 stages. Control plates were inoculated with 500 μL OP50-1. Nematodes were incubated at room temperature for 2 days after infection before collection for RNA extraction. Nematodes were washed off plates with M9 buffer into 15 mL conical tubes, and subjected to gravity floatation 5X to remove excess progeny. Nematodes were concentrated and transferred to microcentrifuge tubes. RNA was extracted using TRI-zol from Direct-zol RNA kit (Zymo Research). cDNA was made from extracted RNA with Maxima H Minus First Strand cDNA Synthesis Kit (Thermo Scientific). qRT-PCR was performed using Brilliant III Ultra-Fast SYBR green QPCR Master Mix (Agilent) on a Stratagene Mx3005P QPCR instrument. Each qRT-PCR was measured in technical duplicate. Gene expression data was normalized to *snb-1* gene expression. ΔΔCt method was used to calculate gene expression fold changes and analysis was performed in Graphpad Prism v10. One-sample t-tests were performed to compare gene expression fold changes to a theoretical mean of 1.

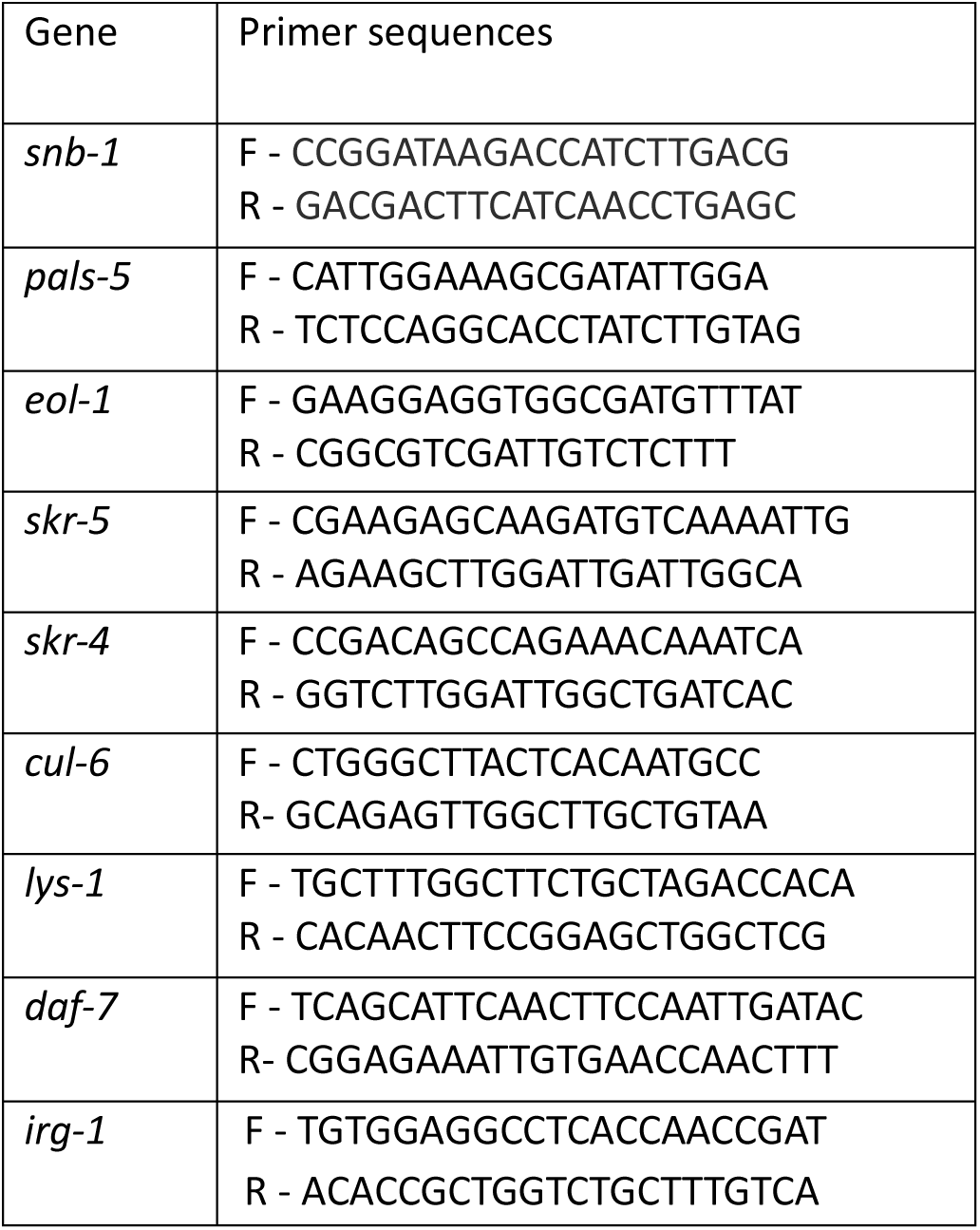

---

**Figure S1:**
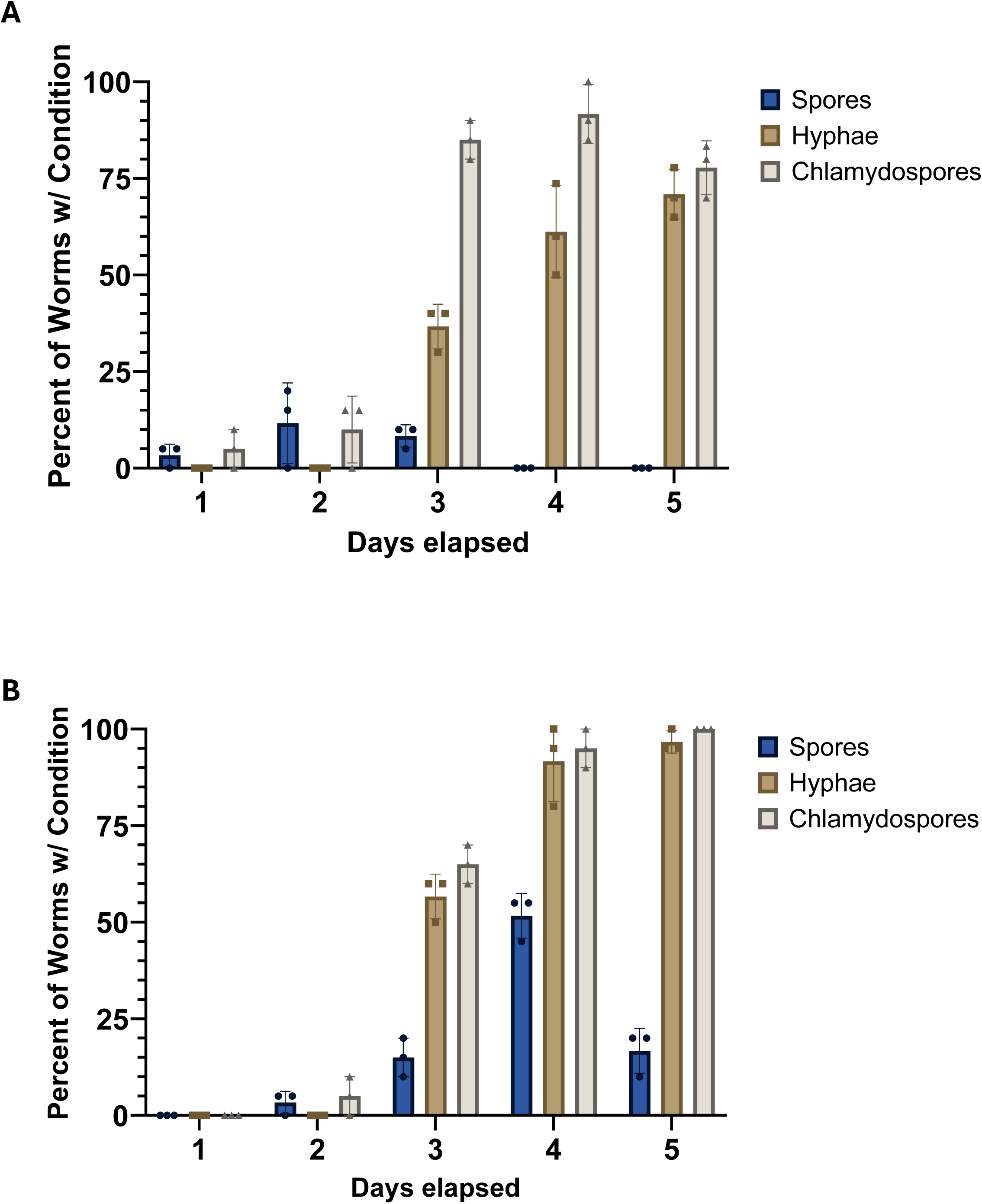
Ingested *M. hiemalis* spores germinate hyphal bodies and accumulate inside of the *Caenorhabditis* nematode intestine. (A) Fluorescence micrographs of *M. hiemalis* growth stages stained with DY96 in infected wild type N2 *C. elegans*. *C. elegans* were scored for fungal growth conditions in biological triplicates. Graphs show the percentage of nematodes with *M. hiemalis* spores, hyphae, or chlamydospores. *M. hiemalis* growth stage counts in N2 *C. elegans* across 5 days of observation by fluorescence microscopy with (A) adult day 2 (B) adult day 3 nematodes. Experiment was repeated 3 times independently (n = 20 for each experiment). Error bars show standard deviation.

**Figure S2:**
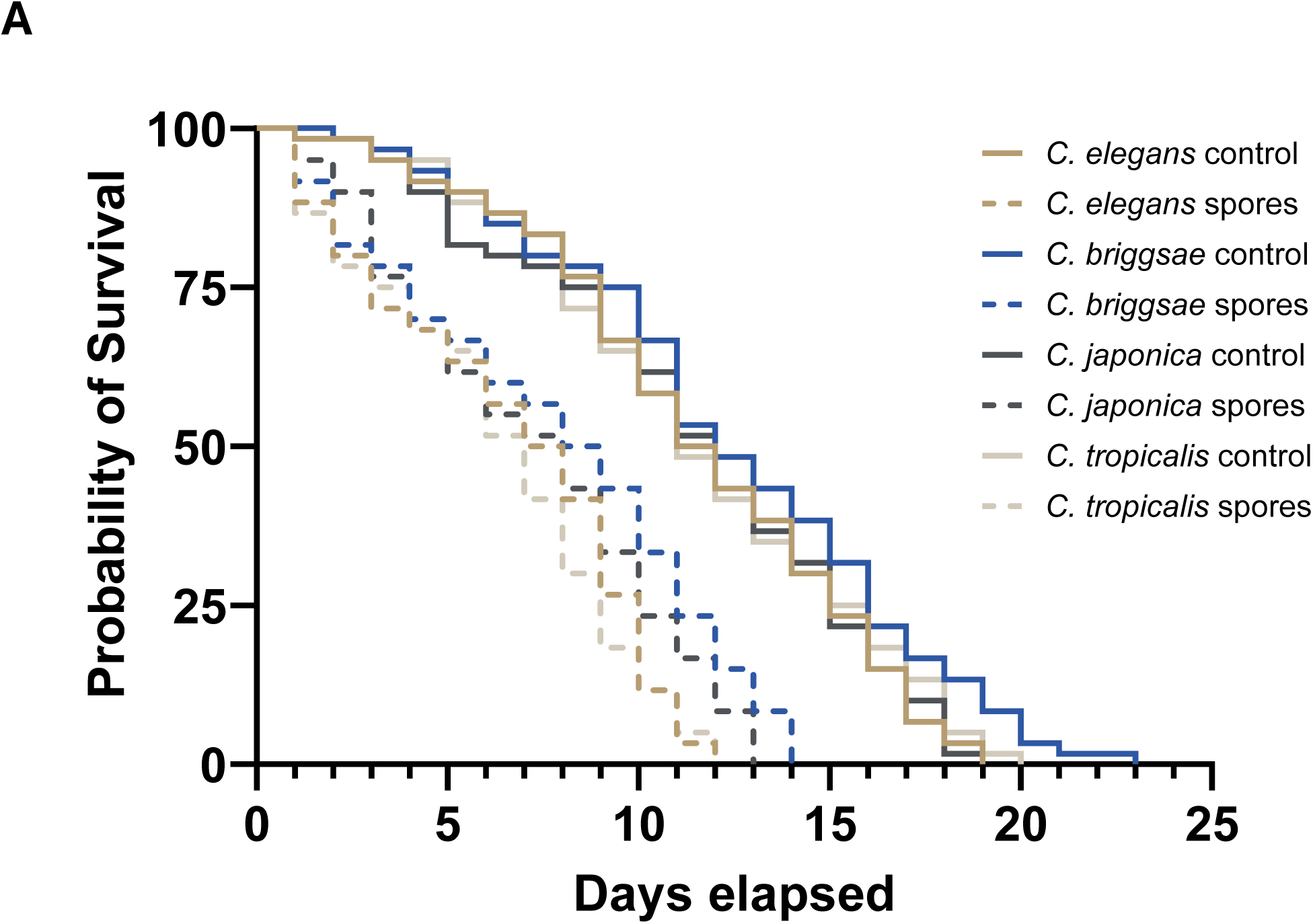
*M. hiemalis* kills *Caenorhabditis* nematodes with a wide host range. (A) Combined survival curves of *Caenorhabditis* strains *C. elegans* (N2), *C. japonica* (DF5081), *C. tropicalis* (JU1373), and *C. briggsae* (wild isolate) grown on *M. hiemalis* infection plates or OP50-1 controls. Nematodes were scored for survival every day for their lifespan. Graphs represent lifespans of biological triplicate groups (n = 20).

